# Dynamic Regulation of Cholesterol Metabolism Genes in Dopaminergic Neurons following Methamphetamine Treatment as Revealed by Single-Nucleus RNA Sequencing

**DOI:** 10.1101/2025.07.28.667272

**Authors:** Wei Sun, Yajun Zhang, Snehashis Roy, Sarah K. Williams Avram, Ching-Yu Sun, Timothy J. Petros, Susan G. Amara

## Abstract

Methamphetamine profoundly alters the function of midbrain dopaminergic neurons, yet the molecular mechanisms underlying these effects are not fully understood. Using single-nucleus RNA sequencing, we found that acute methamphetamine exposure leads to a marked up-regulation of cholesterol metabolism genes in dopaminergic neurons of the ventral tegmental area and substantia nigra—a response that was much less pronounced in astrocytes and largely absent in other cell types. Further analysis using a ribosome tagging strategy combined with RNA sequencing revealed that amphetamine, a structurally similar psychostimulant, induced similar gene expression changes, whereas methylphenidate, a structurally distinct psychostimulant, did not, highlighting drug-specific transcriptional responses. Notably, repeated methamphetamine exposure resulted in down-regulation of cholesterol metabolism genes in dopaminergic neurons. Interestingly, we also observed that, compared to neighboring cell types, dopaminergic neurons are highly enriched in genes encoding cholesterol biosynthesis enzymes, including the rate-limiting enzyme *Hmgcr*, and key regulators *Srebf2* and *Insig1*, challenging the prevailing view that neurons rely mainly on astrocyte-derived cholesterol. In summary, our study highlights dynamic changes in cholesterol metabolism in dopaminergic neurons in response to amphetamines and uncovers the potential importance of cholesterol homeostasis for dopaminergic neuron function.

## Introduction

Methamphetamine is a potent psychostimulant that profoundly affects the central nervous system, primarily by targeting dopaminergic neurons. Mechanistically, methamphetamine binds to the dopamine transporter (DAT), inhibits reuptake, induces subsequent efflux of dopamine and elevates extracellular dopamine concentrations (Amara and Sonders 1998). Previous work from our lab and others has shown that methamphetamine and amphetamine also activate an intracellular G-protein coupled receptor, TAAR1, and several downstream signaling pathways, resulting in activation of protein kinase A (PKA), and the small GTPases, RhoA and Rac1 (Bunzow, Sonders et al. 2001, Wheeler, Underhill et al. 2015, Underhill, Hullihen et al. 2021). The activation of these intracellular signaling pathways by methamphetamine has the potential to alter gene expression patterns in dopaminergic neurons.

The majority of dopaminergic neurons reside in the midbrain, specifically in the ventral tegmental area (VTA) and the substantia nigra pars compacta (SNc). Dopaminergic neurons in the VTA project primarily to the nucleus accumbens, prefrontal cortex, and limbic regions, playing a central role in the brain’s reward circuitry and addiction. In contrast, dopaminergic neurons in the SNc project to the dorsal striatum and are essential for the regulation of voluntary movement and motor control. Most previous genomics and proteomics studies have focused on structures downstream of dopaminergic neurons, such as the hippocampus and cortex (Freeman, Brebner et al. 2005, Zeng, Yu et al. 2023, Li, Yu et al. 2024, Oladapo, Deshetty et al. 2025). However, whether methamphetamine alters gene expression in dopaminergic neurons themselves remains unknown.

Recently, single-nucleus RNA-seq has emerged as a powerful tool for studying transcriptional changes in individual cell types, including dopaminergic neurons, in both drug addiction and neurodegeneration (Agarwal, Sandor et al. 2020, Kilfeather, Khoo et al. 2024). This approach offers high resolution and avoids biases introduced by tissue dissociation, enabling analysis of rare or vulnerable neuronal populations as well as glial cells.

In this study, we used single-nucleus RNA-sequencing to directly investigate gene expression changes in dopaminergic neurons of the VTA and SNc *in vivo* following acute methamphetamine treatment. This approach allowed us to assess both global transcriptional changes in dopaminergic neurons and responses in neighboring cell types. We found striking up-regulation of cholesterol metabolism genes in dopaminergic neurons, including key enzymes, transcription factors, and regulatory genes. While astrocytes also upregulated some cholesterol metabolism genes, the response was less broad and robust than in dopaminergic neurons. No other major cell types exhibited such changes. Interestingly, our findings reveal that dopaminergic neurons are highly enriched for genes involved in cholesterol metabolism, challenging the longstanding view that neurons rely primarily on astrocytes for cholesterol supply.

In addition to their potential for abuse, psychostimulants are commonly prescribed to treat symptoms of ADHD, with amphetamines and methylphenidate being two of the most widely used medications. To examine the effects of amphetamine and methylphenidate on gene expression specifically in dopaminergic neurons, we performed bulk RNA-seq using tyrosine hydroxylase RiboTag mice (*Th-cre:RiboTag*). The mouse line enables selective immunoprecipitation and isolation of ribosome-bound mRNAs from dopaminergic neurons. We found that amphetamine, like methamphetamine, induced up-regulation of cholesterol metabolism genes, whereas methylphenidate did not produce this effect, underscoring divergent mechanisms and effects between amphetamines, which enter the cell, and methylphenidate, which binds to the DAT and blocks dopamine uptake.

## Results

### Single-nucleus RNA sequencing of the VTA and SNc

To understand the impact of acute methamphetamine injection on gene expression within midbrain dopaminergic neurons and their surrounding cells, we performed single-nucleus RNA sequencing (snRNA-seq) using Dat bacTrap mice. In this mouse line, ribosomes within dopaminergic neurons are genetically labeled with EGFP, enabling precise microdissection of the VTA and SNc under fluorescence guidance, thereby minimizing contamination from adjacent brain regions (Fig. 1A). Previous studies have also demonstrated the utility of this mouse line for sorting nuclei and transcriptome studies (Yaghmaeian Salmani, Lahti et al. 2024).

**Figure 1.**
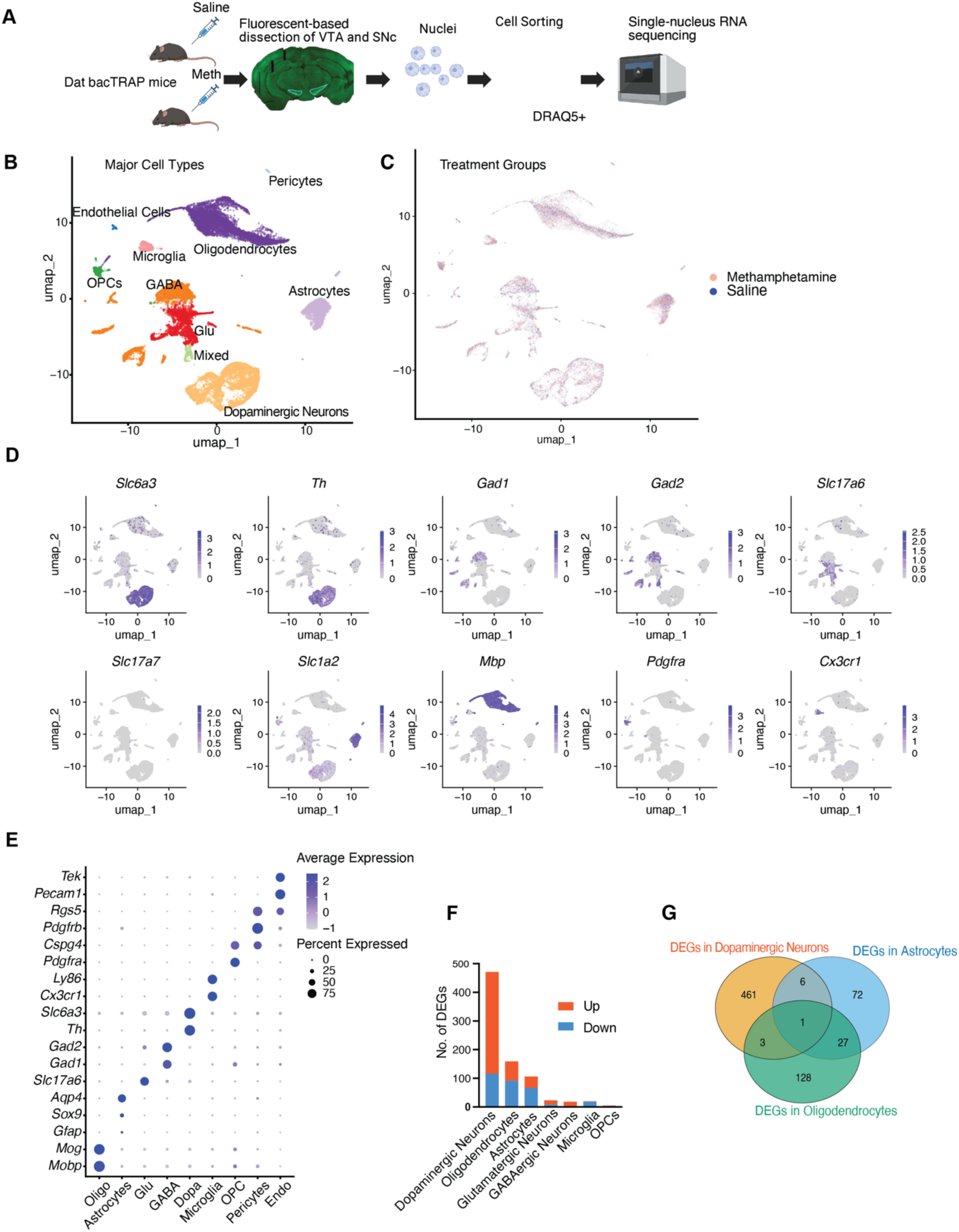
Overview of snRNA-seq experimental design and data analysis. (A) Schematic outlining the experimental workflow. *Dat bacTRAP* mice were injected intraperitoneally with either saline or methamphetamine (3 mg/kg). Tissue from the VTA and SNc was dissected 3 hours post-injection based on EGFP fluorescence. Isolated nuclei were labeled with DRAQ5 and sorted for single-nucleus RNA sequencing. (B) UMAP visualization of all nuclei, showing the major cell types identified. (C) UMAP plot showing the distribution of nuclei from methamphetamine- and saline-treated mice. (D) Dot plot depicting expression of canonical marker genes across major cell types. (E) Feature plots showing expression of selected marker genes projected onto the UMAP embedding. (F) Numbers of differentially expressed genes (DEGs) between saline- and methamphetamine-treated mice (FDR < 0.05, fold change > 1.3), shown for each major cell type with >500 nuclei detected. (G) Venn diagram illustrating overlap of DEGs among dopaminergic neurons, astrocytes, and oligodendrocytes. **Abbreviations:** OPC, oligodendrocyte precursor cells; GABA, GABAergic neurons; Glu, glutamatergic neurons; Dopa, dopaminergic neurons; Oligo, oligodendrocytes; Endo, endothelial cells; Mixed, neurons expressing multiple markers.

A total of 16 mice (4 males and 4 females per condition) aged 8 to 16 weeks were used. Mice received either saline or methamphetamine, and tissue was harvested 3 hours post-injection for nuclei preparation. Following FACS enrichment for EGFP^+^ nuclei, EGFP^-^ nuclei were also subsequently collected to increase cell numbers and provide transcriptomic insights in non-dopaminergic neurons (Supplementary Fig.1A). snRNA-seq was performed using the 10X Genomics Chromium v3 platform. After filtering of low-quality nuclei, we retained 29,539 high-quality nuclei for downstream analysis, including 13,191 nuclei from saline-treated mice and 16,348 from methamphetamine-treated mice.

Following batch correction, unbiased clustering and UMAP visualization revealed comparable cell type distributions across saline and methamphetamine-treated samples, with no evidence of major cell state transitions or gross compositional shifts after acute methamphetamine exposure (Fig. 1B, C). This indicates that, at this early time point, methamphetamine does not induce widespread changes in cell identity or population structure within the midbrain.

Cell types were annotated and verified based on the expression of canonical marker genes: dopaminergic neurons (*Slc6a3, Th*), GABAergic interneurons (*Gad1, Gad2*), glutamatergic neurons (*Slc17a7*), astrocytes (*Slc1a4, Gfap, Sox9, Aqp4*), oligodendrocytes (*Mbp, Mog, Mobp*), oligodendrocyte precursor cells (*Pdgfra, Cspg4*), microglia (*Cx3cr1, Ly86*), pericytes (*Pdgfrb, Rgs5*), and endothelial cells (*Tek, Pecam1*) (Fig.1D, E). These cell types are consistent with prior single cell studies of the VTA and/or SNc (Agarwal, Sandor et al. 2020, Phillips, Tuscher et al. 2022, Wu, Zhang et al. 2024, Yaghmaeian Salmani, Lahti et al. 2024).

5,587 nuclei were identified as dopaminergic neurons, with the remaining nuclei representing other midbrain cell types. Oligodendrocytes constituted the most populous cell type in the data, followed by dopaminergic neurons, GABAergic neurons, and astrocytes. Similar numbers of nuclei from saline and methamphetamine-treated groups were detected for each cell type (Supplementary Fig.1B). Notably, RNA counts per cell detected in neurons were higher than in glial cells (Supplementary Fig. 1C). This is likely a reflection of the difference in total RNA content in neuronal and non-neuronal cell types.

Consistent with previous studies, VTA and SNc glutamatergic neurons predominantly expressed vGlut2 (*Slc17a6*) (Yamaguchi, Sheen and Morales 2007), but not vGlut1 (*Slc17a7*). We also identified a distinct population co-expressing the glutamatergic marker *Slc17a6* and the GABAergic marker *Gad2* (Fig.1B, D) (referred to as “mixed neurons” in this study), potentially reflecting GABA-glutamate co-transmitting neurons (Ntamati and Luscher 2016).

### Differential Gene Expression in Response to Methamphetamine Across Major Cell Types

Next, we examined differentially expressed genes (DEGs) in major cell types between saline and methamphetamine-treated animals. We found that dopaminergic neurons exhibited the highest number of DEGs at a significance level of FDR < 0.05 and an absolute fold change > 1.3, with 355 genes up-regulated and 116 genes down-regulated. This was followed by oligodendrocytes, which had 67 genes up-regulated and 92 genes down-regulated, and astrocytes, with 40 genes up-regulated and 66 genes down-regulated. In contrast, other cell types showed significantly fewer DEGs, including glutamatergic neurons (23 DEGs), GABAergic neurons (18 DEGs), microglia (20 DEGs), and oligodendrocyte precursor cells (OPCs) (5 DEGs). It is well-known that dopaminergic neurons are major targets of methamphetamine, and our results reinforce the importance of these neurons in the acute response to methamphetamine. It is notable that GABAergic and glutamatergic neurons in the VTA exhibited far fewer DEGs. On the other hand, oligodendrocytes and astrocytes showed more DEGs than these neurons (Fig. 1F).

We then examined the overlap of DEGs among the cell types that exhibited the most changes. The majority of DEGs were unique to each cell type. There was greater overlap between astrocytes and oligodendrocytes (28 genes) than between dopaminergic neurons and glia (7 genes between dopaminergic neurons and astrocytes, and 4 genes between dopaminergic neurons and oligodendrocytes) (Fig.1G). These data suggest that each cell type has unique mechanisms in responding to methamphetamine, with some similarities between oligodendrocytes and astrocytes.

### Up-regulation of genes involved in cholesterol metabolism in dopaminergic neurons

Dopaminergic neurons play a pivotal role in the brain’s response to methamphetamine. Methamphetamine increases dopamine release and inhibits its reuptake by blocking DAT, leading to elevated dopamine levels in the synaptic cleft and prolonged signaling (Fleckenstein, Volz et al. 2007). Our findings indicate that following acute methamphetamine treatment, dopaminergic neurons exhibit the most pronounced changes in gene expression. Notably, the top 10 most upregulated genes are all associated with lipid metabolism. 8 of these genes (*Lss, Ldlr, Mvd, Msmo1, Nsdhl, Fdps, Acat2, Hmgcs1*) are involved in cholesterol metabolism, while the remaining two genes (*Fads2, Scd2*) are related to fatty acid metabolism (Fig.2A). Metascape enrichment analysis (Zhou, Zhou et al. 2019) further identified “Cholesterol metabolism with Bloch and Kandutsch-Russell Pathways” as the top pathway (Fig.2B).

**Figure 2.**
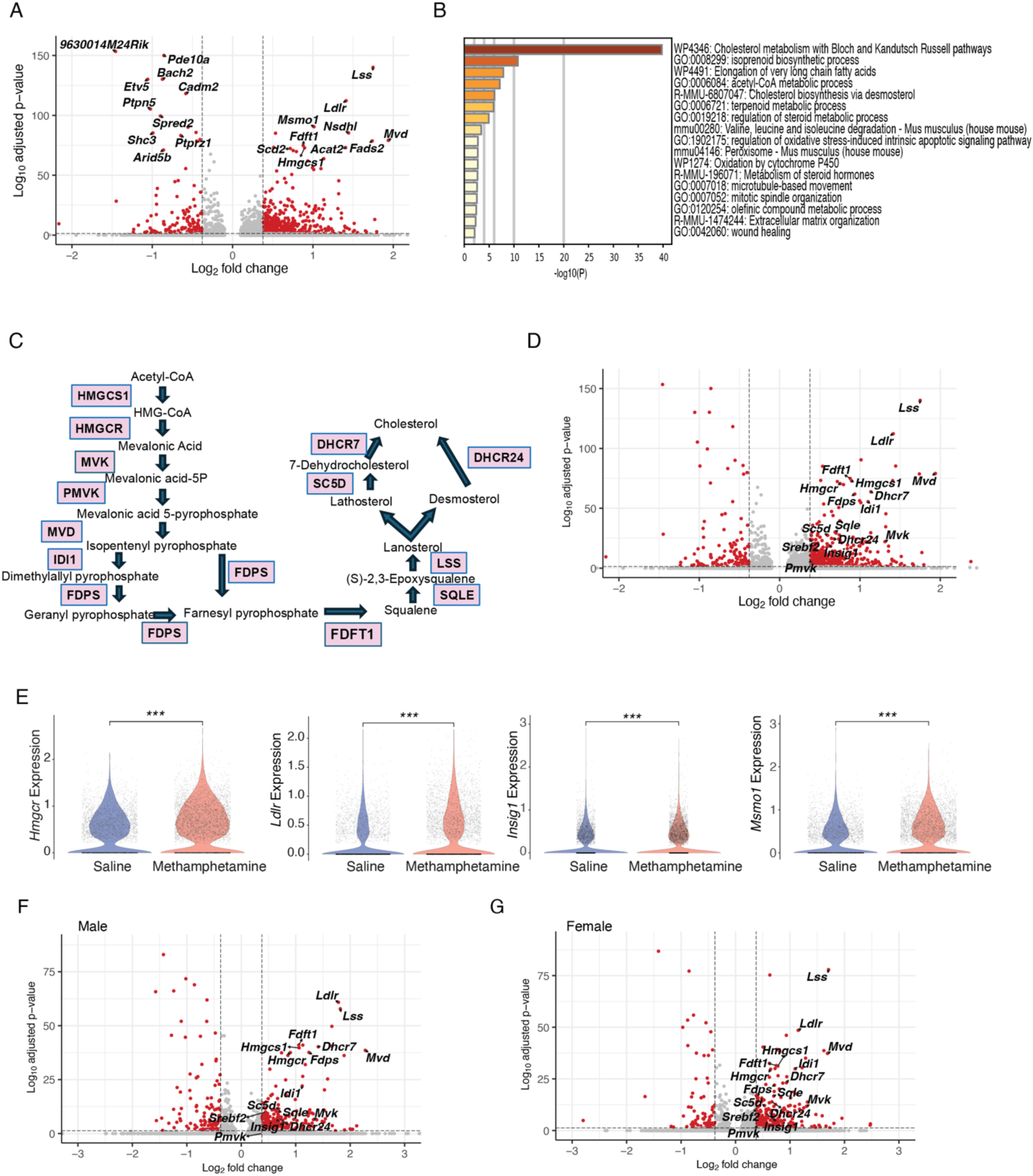
Differential gene expression and pathway analysis in dopaminergic neurons following acute methamphetamine treatment. (A) Volcano plot displaying the top 10 upregulated and downregulated genes (by adjusted p-value) in dopaminergic neurons after acute methamphetamine administration. Red dots indicate genes with a false discovery rate (FDR) < 0.1 and absolute fold change > 1.3. (B) Metascape functional enrichment analysis highlighting the top upregulated pathways in dopaminergic neurons. (C) Schematic of key genes in the cholesterol synthesis pathway, adapted from (Sitaula and Burris 2016). (D) Volcano plot showing adjusted p-values and log₂ fold changes for genes encoding enzymes depicted in (C), as well as *Ldlr, Insig1, and Srebf2*. Red dots indicate genes with FDR < 0.1 and absolute fold change > 1.3 when comparing methamphetamine- and saline-treated dopaminergic neurons. (E) Violin plots illustrating expression of representative differentially expressed genes (DEGs) in dopaminergic neurons between methamphetamine and saline groups. ***, p < 0.001, Wilcoxon rank-sum test. (F, G) Volcano plots showing adjusted p-values and log₂ fold changes of cholesterol metabolism–related genes in male (F) and female (G) dopaminergic neurons. Red dots indicate FDR < 0.1 and absolute fold change > 1.3.

Subsequently, we examined the expression of genes encoding key enzymes (*Hmgcs1, Hmgcr, Mvk, Pmvk, Mvd, Idi1, Fdps, Fdft1, Sqle, Lss, Sc5d, Dhcr24, Dhcr7*) in the cholesterol synthesis pathway (Fig.2C) (Sitaula and Burris 2016), as well as genes involved in cholesterol uptake (*Ldlr*) and cholesterol regulation (*Insig1, Srebf2*). All these genes were upregulated following acute methamphetamine treatment in dopaminergic neurons (Fig. 2D, E). Importantly, *Hmgcr*, the gene encoding the rate-limiting enzyme 3-hydroxy-3-methylglutaryl-CoA reductase (HMGCR), was significantly upregulated. Also, the key transcription factor in regulating cholesterol synthesis genes (Eberle, Hegarty et al. 2004), *Srebf2*, is also upregulated, which is in concordance with the broad up-regulation of genes encoding important enzymes for cholesterol synthesis. In summary, our data demonstrated that dopaminergic neurons upregulate cholesterol synthesis, uptake and regulation in response to acute methamphetamine treatment. Interestingly, enzymes specific to both the Bloch (*Dhcr7*) and the Kandutsch-Russell (*Dhcr24*) cholesterol synthesis pathways are both upregulated, although it is commonly believed that the Kandutsch-Russell pathway is the predominant pathway in neurons (Nieweg, Schaller and Pfrieger 2009).

Sex differences in cholesterol metabolism have been well documented by previous studies(Wang, Magkos and Mittendorfer 2011, Holven and Roeters van Lennep 2023). To examine whether methamphetamine-induced differential expression in cholesterol metabolism exists in both males and females, we divided the samples based on *Xist* expression, a gene only expressed in female cells (Supplementary Fig. 2A, B). The validity of this separation was confirmed by examining the expression of *Xist* (female only), *Uty* (male only), and *Kdm5d* (male only) (Supplementary Fig. 2C). Consistently, genes involved in cholesterol metabolism, uptake, and regulation were among the top upregulated genes in both males and females (Fig. 2F, G, Supplementary Fig. 2D).

Dopaminergic neurons in the VTA and SNc are anatomically and functionally distinct. Dopaminergic neurons in the VTA primarily participate in mesolimbic and mesocortical pathways, regulating reward, motivation, emotion, and cognition. In contrast, dopaminergic neurons from the SNc project predominantly to the dorsal striatum, playing a key role in motor control. Previous studies have demonstrated that *Sox6* expression is enriched in SNc dopaminergic neurons, whereas *Calb1* expression is primarily found in the VTA dopaminergic neurons (Poulin, Zou et al. 2014, Yaghmaeian Salmani, Lahti et al. 2024). Consistent with these findings, UMAP visualization of dopaminergic neurons indicated distinct distributions of *Sox6*+ and *Calb1*+ dopaminergic neurons (Supplementary Fig. 3A, B). To evaluate differential responses to acute methamphetamine exposure between VTA and SNc dopaminergic neurons, we categorized the dopaminergic neuron population into four subpopulations: *Sox6*-only, *Calb1*-only, double-positive, and double-negative (Supplementary Fig. 3C). Subsequent analyses revealed significant up-regulation of genes associated with cholesterol metabolism across all four dopaminergic subpopulations (Supplementary Fig. 3D). These results indicate that acute methamphetamine treatment broadly activates cholesterol metabolic pathways in both VTA and SNc dopaminergic neurons.

To verify findings from our single-nucleus RNA-seq study, we performed RNAscope to assess representative genes upregulated following treatment. The selected genes included *Ldlr* (responsible for cholesterol uptake), *Hmgcr* (the rate-limiting enzyme for cholesterol synthesis), *Insig1* (involved in cholesterol regulation), and *Msmo1* (one of the top upregulated genes encoding an enzyme essential for cholesterol synthesis) (Fig.3A, B). Dopaminergic neuron-specific genes *Slc6a3* and *Th* were utilized to define the area occupied by dopaminergic neurons. Quantification of RNAscope dots per area occupied by dopaminergic neurons provided an estimate of mRNA abundance. The VTA was chosen as the area of interest because it is the main region that mediates the downstream effects of methamphetamine. All four genes demonstrated significant up-regulation in VTA dopaminergic neurons (Fig.3C). These results support our transcriptomic findings and highlight a robust activation of cholesterol metabolism pathways within dopaminergic neurons following acute methamphetamine treatment.

**Figure 3.**
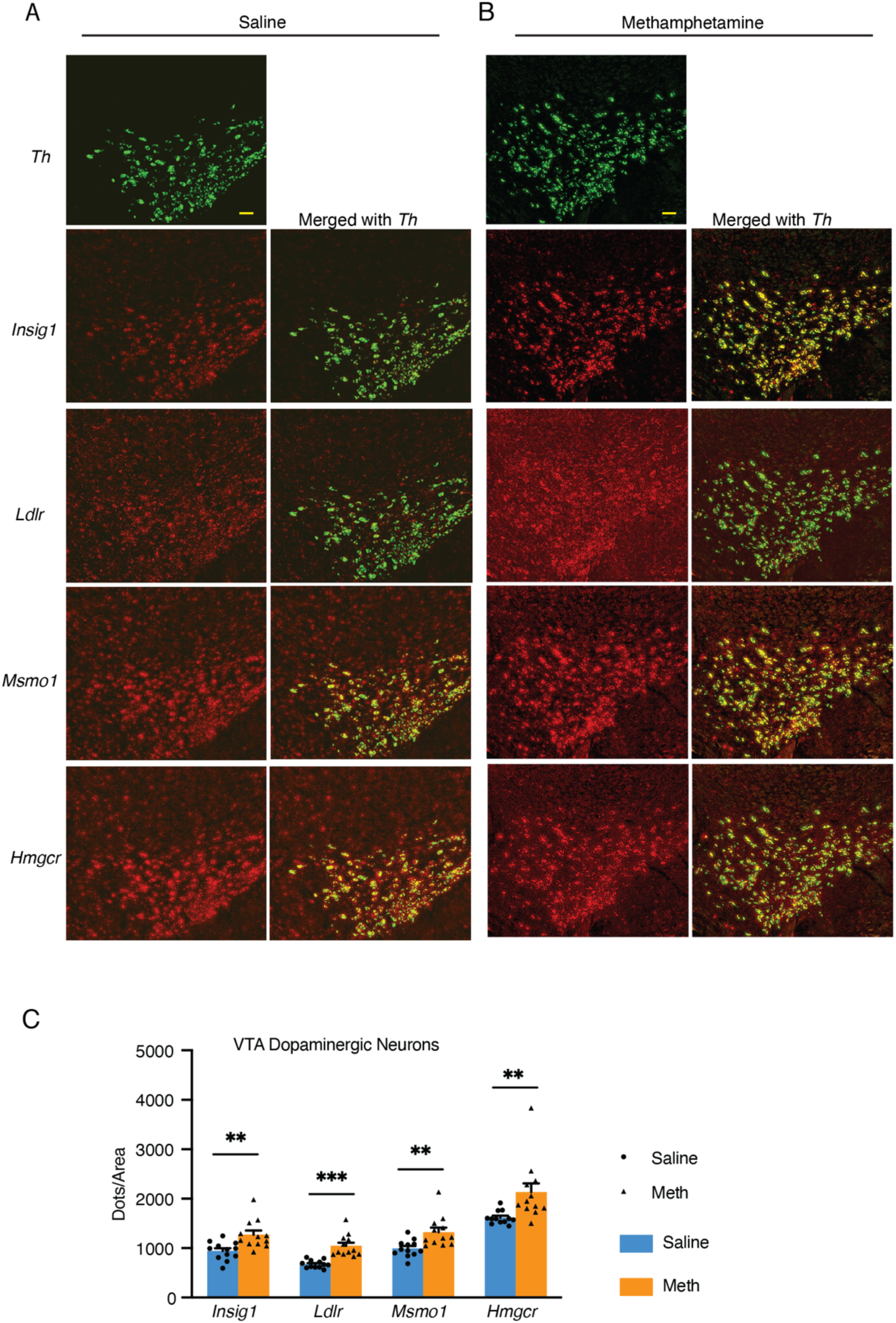
RNAscope analysis of selected cholesterol metabolism genes in dopaminergic neurons following acute methamphetamine treatment. (A, B) Representative RNAscope HiPlex images showing dopaminergic neurons identified by *Th* (green), cholesterol metabolism genes *Insig1* (red), and *Ldlr* (red), *Msmo1*(red), and *Hmgcr* (red) in saline- (A) and methamphetamine- (B) treated mice. Scale bar, 100 μm. (C) Quantification of RNAscope signal for selected genes in dopaminergic neurons from saline- and methamphetamine-treated mice. Expression is represented as dots per ROI. Statistical significance was assessed using the unpaired t-test; *p < 0.05, **p < 0.01, ***p < 0.001, error bar: SEM.

### Up-regulation of Cholesterol Metabolism Genes in Astrocytes But not Oligodendrocytes

Cholesterol is a fundamental component of cell membranes and is vital for synaptic transmission. Cholesterol cannot readily pass the blood-brain barrier. Astrocytes are currently considered the primary source of cholesterol for neurons (Li, Zhang and Liu 2022). Thus, we analyzed differential gene expression in astrocytes following acute methamphetamine treatment in the VTA and SNc. Pathway analysis using WikiPathways 2024 (Agrawal, Balci et al. 2024) revealed that 11 out of the 17 upregulated pathways were related to cholesterol and lipid metabolism, including “Sterol Regulatory Element Binding Proteins (SREBP) signaling”, “SREBF and miR-33 in Cholesterol and Lipid Homeostasis”, and “Cholesterol metabolism” (Fig.4A). Key differentially expressed genes (DEGs) in these pathways included: *Srebf1*, encoding SREBP-1 (Sterol Regulatory

**Figure 4.**
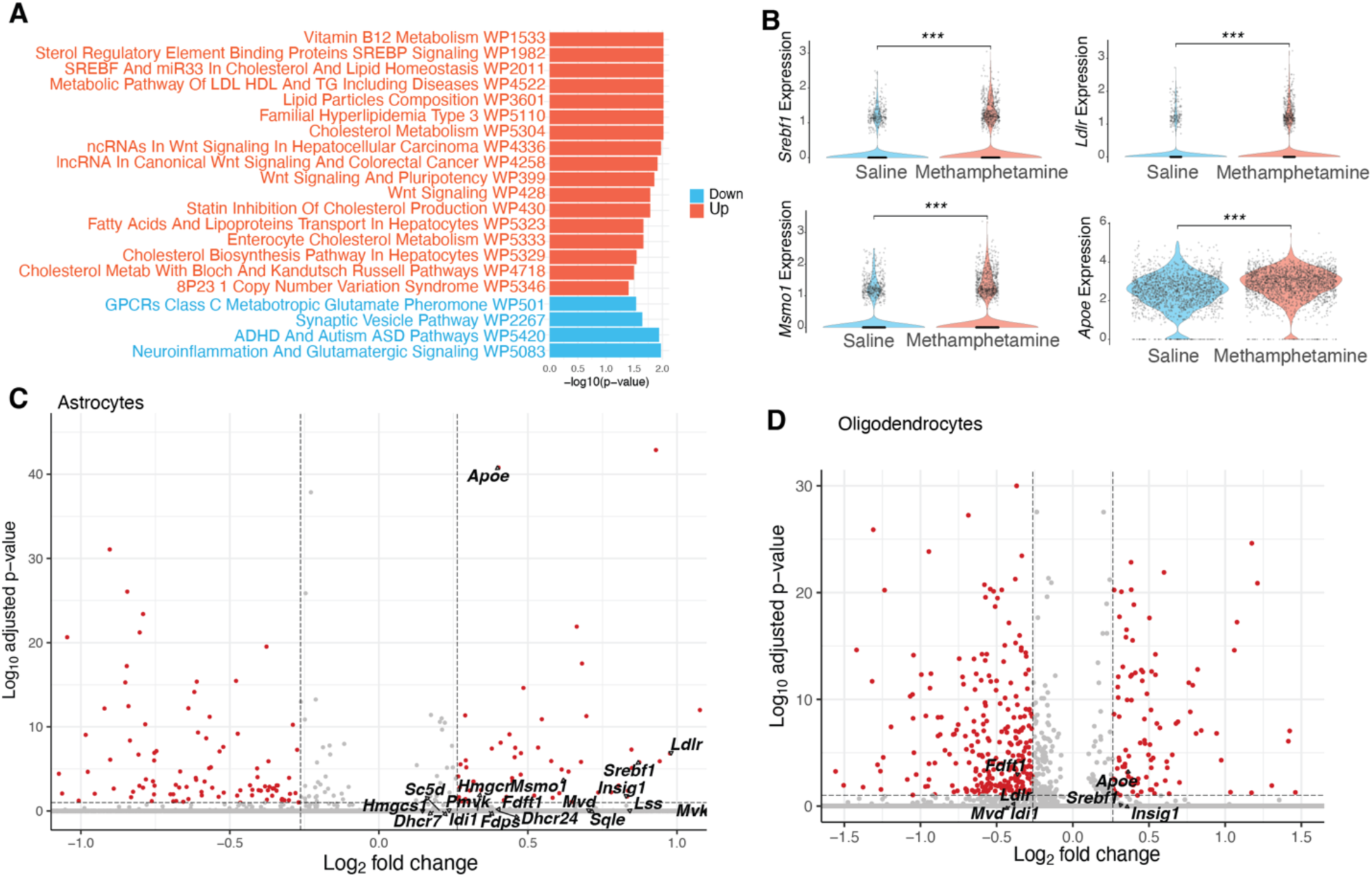
Cholesterol metabolism genes in astrocytes and oligodendrocytes. (A) Pathway analysis of differentially expressed genes (DEGs) in astrocytes induced by acute methamphetamine treatment, based on WikiPathways. (B) Violin plots showing expression levels of significant DEGs in the SREBP pathway. For each gene, saline and methamphetamine groups were compared using the Wilcoxon rank-sum test; *** p < 0.001. (C, D) Volcano plots showing adjusted p-values and log_2_ fold changes for genes associated with cholesterol metabolism in astrocytes (C) and oligodendrocytes (D) (cutoff: FDR < 0.1, absolute fold change > 1.2).

Element-Binding Protein-1), a major transcription factor regulating lipid metabolism; *Ldlr,* which encodes the receptor for extracellular cholesterol uptake; *Msmo1*, encoding an enzyme involved in cholesterol biosynthesis; *Apoe*, encoding ApoE lipoprotein, essential for transporting cholesterol and phospholipids from astrocytes to neurons (Fig.4B, C). These results suggested that astrocytes also upregulate cholesterol synthesis and transport in response to acute methamphetamine treatment, possibly providing more cholesterol to neurons. The up-regulation of cholesterol metabolism genes suggests a possible disruption in lipid homeostasis by acute methamphetamine treatment.

Cholesterol metabolism is essential in oligodendrocytes, as cholesterol constitutes a major component of myelin and is required for both developmental myelination and remyelination after injury (Saher, Brügger et al. 2005). We examined whether genes involved in cholesterol metabolism are altered in oligodendrocytes following acute methamphetamine treatment. Surprisingly, most of these genes were below the detection threshold in oligodendrocytes and, therefore, could not be visualized in the volcano plot. Of the 18 genes examined, only 7 were detected in oligodendrocytes, and only two of these showed significant changes, but in opposite directions: *Fdft1*, which encodes an enzyme in cholesterol synthesis, was downregulated, while *Apoe* was upregulated (Fig.4D). Thus, unlike in dopaminergic neurons and astrocytes, our data did not indicate major disruptions in cholesterol metabolism in oligodendrocytes. In addition, excitatory and inhibitory neurons also did not show major changes in cholesterol-related genes (Supplementary Fig.5A, B).

### Dopaminergic neurons are highly enriched in genes involved in cholesterol metabolism

We observed up-regulation of cholesterol metabolism genes in both dopaminergic neurons and astrocytes. Although astrocytes have traditionally been considered the primary source of cholesterol for neurons, our RNAscope data revealed that *Hmgcr*—the gene encoding the rate-limiting enzyme in cholesterol biosynthesis—along with *Msmo1* (an enzyme in cholesterol metabolism) and *Insig1* (a regulator of cholesterol metabolism), were highly expressed in dopaminergic neurons of both the VTA and SNc (Fig. 3A, B; Fig. 5A-E). Supporting this, *in situ* hybridization images from the Allen Brain Atlas also indicate enrichment of these genes in both regions, consistent with a dopaminergic distribution (Supplementary Fig. 4A-D), which significantly differs from patterns of genes with an astrocytic enrichment, such as *Apoe* (Supplementary Fig. 4E).

**Figure 5.**
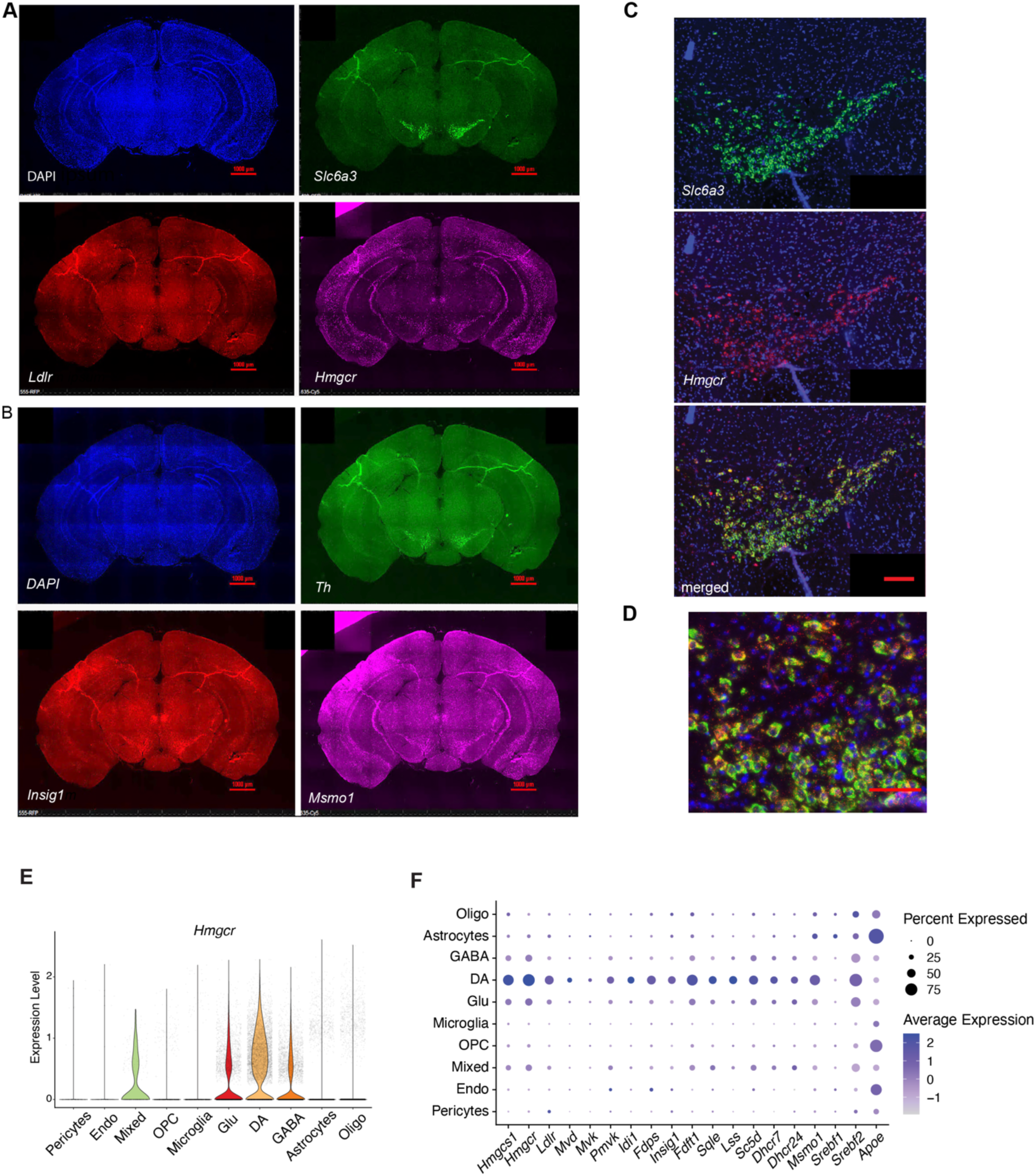
RNAscope analysis of selected cholesterol metabolism genes in dopaminergic neurons. (A) Representative RNAscope HiPlex images showing DAPI-labeled nuclei (blue), dopaminergic neurons identified by *Slc6a3* (green), *Ldlr* (red), and *Hmgcr* (purple). Scale bar, 1,000 μm. (B) Representative RNAscope HiPlex images on the same brain section as in (A) with DAPI (blue), *Th* (green) labeling dopaminergic neurons, *Insig1* (red), and *Msmo1* (purple). Scale bar, 1,000 μm. (C) Representative image of RNAscope co-labeling for *Slc6a3* (green) and *Hmgcr* (red). Scale bar, 200 μm. (D) Co-labeling of *Slc6a3* (green) and *Hmgcr* (red). The region occupied by *Slc6a3*-positive cells was defined as the region of interest (ROI), and red dots within the ROI were quantified for *Hmgcr* expression (Scale bar, 50 μm). (E) Violin plot showing *Hmgcr* expression in different cell types in the VTA and SNc. (F) Dot plot depicting expression levels of cholesterol metabolism-associated genes across different cell types. **Abbreviations:** OPC, oligodendrocyte precursor cells; GABA, GABAergic neurons; Glu, glutamatergic neurons; DA, dopaminergic neurons; Oligo, oligodendrocytes; Endo, endothelial cells; Mixed, neurons expressing multiple markers.

To further investigate, we examined *Hmgcr* expression across all major cell types in the VTA and SNc and found that dopaminergic neurons exhibited marked enrichment of *Hmgcr* compared to other cell types (Fig. 5E). Extending this analysis to 18 genes involved in cholesterol metabolism, we found that all genes encoding major cholesterol synthesis enzymes—as well as *Srebf2* (encoding the major transcription factor regulating cholesterol synthesis genes), *Ldlr* (involved in cholesterol uptake) and *Insig1*—were significantly enriched in dopaminergic neurons but not in astrocytes. In contrast, astrocytes showed higher expression of the transcription factor *Srebf1* and the lipoprotein *Apoe* (Fig. 5F).

Contrary to the prevailing view that neurons are dependent on astrocyte-derived cholesterol, our data suggest that dopaminergic neurons possess a robust intrinsic capacity for cholesterol biosynthesis and may be less reliant on astrocytes for cholesterol supply.

### Acute amphetamine treatment upregulates cholesterol metabolism genes, but methylphenidate does not

Amphetamine and methylphenidate are both psychostimulants widely used to treat ADHD. Amphetamine is structurally and functionally similar to methamphetamine; they both increase synaptic dopamine levels by promoting release of dopamine and competing with dopamine as a substrate for DAT. Methylphenidate is structurally different from the amphetamines and works by blocking DAT but with less effect in promoting dopamine release (Challman and Lipsky 2000, Faraone 2018). To determine whether these medications also regulate cholesterol metabolism in dopaminergic neurons, we performed RiboTag RNA-seq using *Th-cre:RiboTag* mice, which enable immunoprecipitation of ribosome-associated RNA specifically from dopaminergic neurons (Fig. 6A). RNA-seq profiling confirmed enrichment of dopaminergic markers (*Slc18a2, Slc6a3*, *Th*, and *Ddc*) and depletion of markers from other cell types (Fig. 6B).

**Figure 6.**
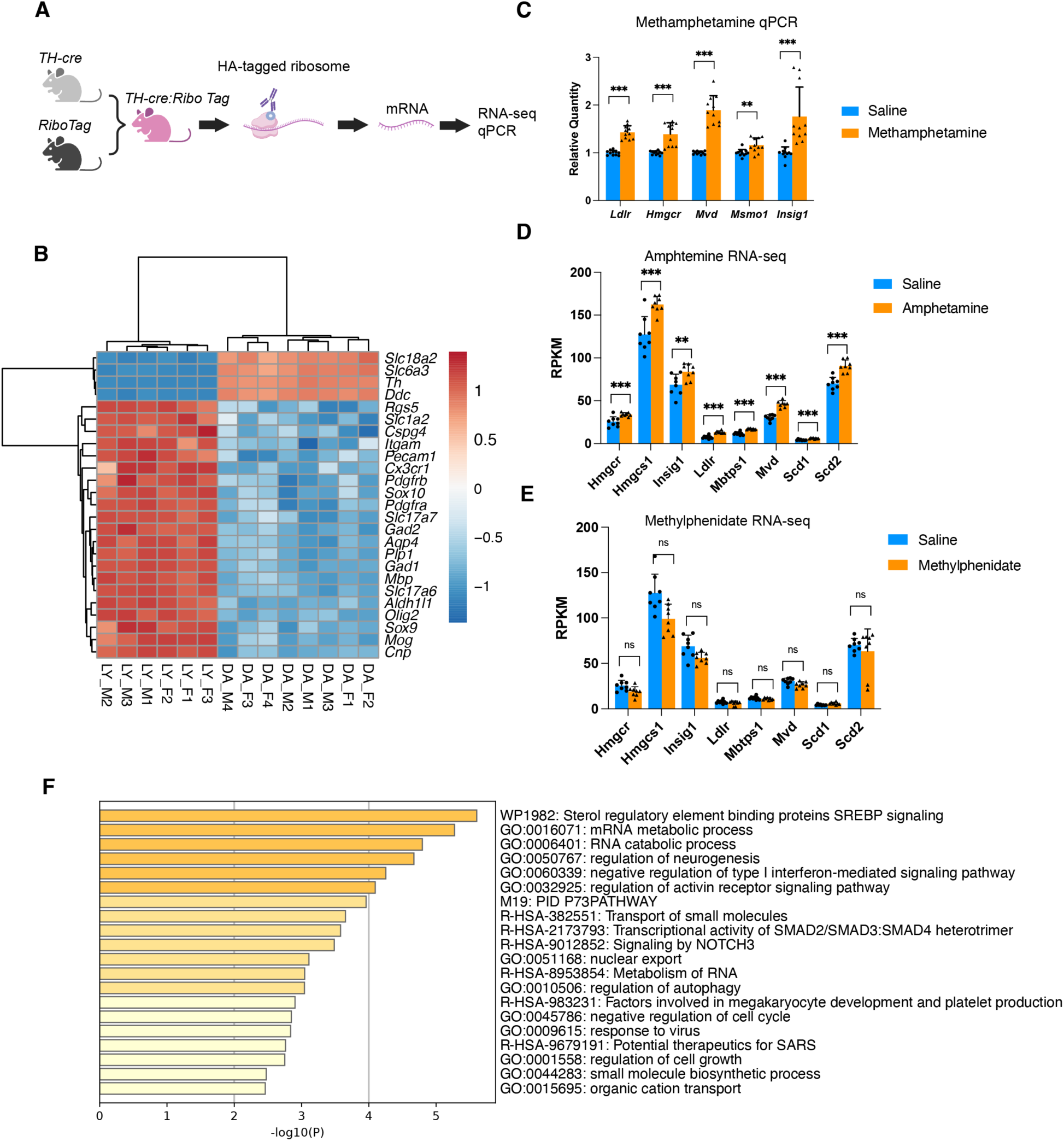
Cholesterol metabolism genes in dopaminergic neurons following acute amphetamine and methylphenidate treatment. (A) Schematic illustrating the *Th-cre:RiboTag* mouse approach used to isolate bulk RNA from dopaminergic neurons for qPCR and RNA-seq analysis. (B) Expression of marker genes in RNA isolated using the RiboTag RNA-seq method. LY, tissue lysate; DA, dopaminergic neurons; F, female; M, male. (C) qPCR validation of cholesterol metabolism–associated genes in RiboTag-isolated RNA from dopaminergic neurons following acute saline or methamphetamine administration. Statistical significance was assessed by unpaired t-test with two-stage step-up FDR correction; **q < 0.01, ***q < 0.001; error bar: standard deviation. (D, E) RNA-seq analysis of genes in the SREBP signaling pathway associated with cholesterol metabolism in dopaminergic neurons after acute amphetamine (D) and methylphenidate (E) treatment in *Th-cre:RiboTag* mice. Statistical significance was determined by unpaired t-test with two-stage step-up FDR correction; **q < 0.01, ***q < 0.001, ns = not significant; error bar: standard deviation. (F) Metascape functional enrichment analysis of upregulated differentially expressed genes (DEGs) in dopaminergic neurons from RiboTag RNA-seq following acute amphetamine treatment.

In mice acutely treated with amphetamine, Metascape functional enrichment analysis of upregulated genes revealed that the top pathway was “Sterol regulatory element binding proteins (SREBP) signaling” (Fig. 6F). Specifically, several genes related to cholesterol metabolism were significantly upregulated, including *Hmgcr, Hmgcs1*, and *Mvd*, which encode the enzymes for cholesterol synthesis; *Mbtps1*, which encodes a key regulator of SREBP-mediated transcription of cholesterol synthesis genes; as well as *Scd1* and *Scd2*, which are targets of the SREBP pathway involved in fatty acid synthesis (Fig. 6D). In contrast, acute methylphenidate treatment did not upregulate any of these genes in dopaminergic neurons (Fig. 6E). Furthermore, using the RiboTag technique, combined with qPCR, we were able to confirm that acute methamphetamine treatment induced genes involved in cholesterol metabolism in dopaminergic neurons similar to those observed in single-nuclei RNA-seq (Fig. 6C).

These results indicate that dopaminergic neurons upregulate cholesterol metabolism genes in response to acute amphetamine and methamphetamine treatment, whereas methylphenidate has no such effect. This is consistent with the structural and mechanistic similarities between amphetamine and methamphetamine, in contrast to the more distinct mechanism of methylphenidate.

### Gene Expression Changes Following Sub-Chronic Methamphetamine Treatment

To further investigate how repeated methamphetamine exposure affects cells in the VTA and SNc, we performed single-nucleus RNA sequencing (snRNA-seq) on mice (16 in total, 4 males and 4 females per treatment group) that received daily injections of methamphetamine or saline for 7 days. Nuclei were isolated from the VTA and SNc 24 hours after the final injection to minimize detection of acute effects. Comparable cell populations were identified across groups, as shown in the UMAP plot (Fig. 7A).

**Figure 7.**
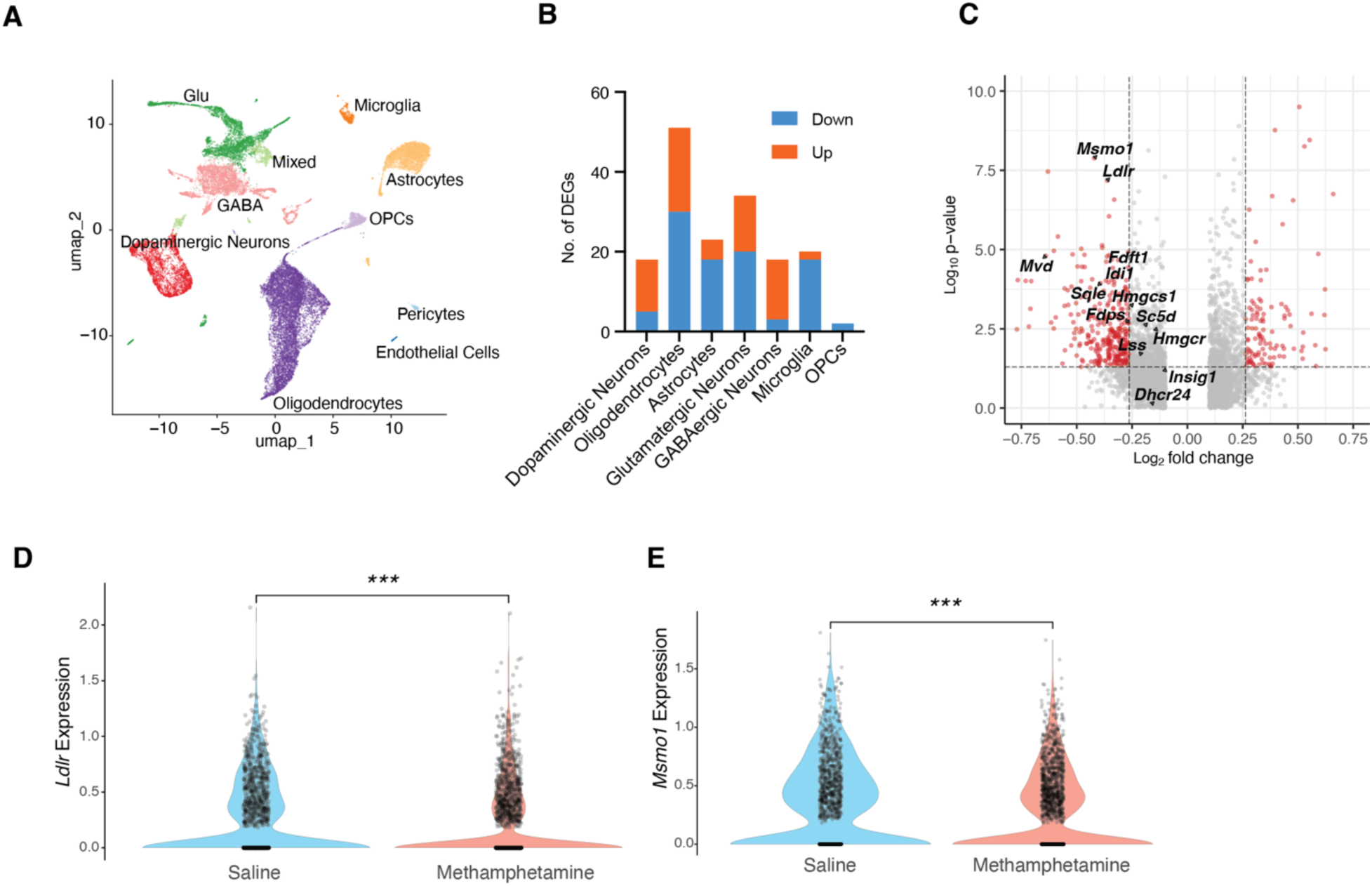
Single-nucleus RNA sequencing of the VTA and SNc following subchronic methamphetamine treatment. (A) UMAP plot showing the major cell populations identified in the VTA and SNc. (B) Numbers of upregulated and downregulated genes detected after 7 days of daily methamphetamine administration (3 mg/kg, intraperitoneally). Differentially expressed genes were identified at FDR < 0.05 and absolute fold change > 1.3. (C) Volcano plot displaying p-values and log₂ fold changes for genes associated with cholesterol metabolism (cutoff: p < 0.05, absolute fold change > 1.2). (D, E) Violin plots comparing *Ldlr* and *Msmo1* expression in dopaminergic neurons from repeated saline- and methamphetamine-treated mice. Statistical significance was assessed by Wilcoxon rank-sum test; ***p < 0.001.

In contrast to the acute methamphetamine treatment—where dopaminergic neurons exhibited the most differentially expressed genes (DEGs)—the sub-chronic methamphetamine group showed the greatest transcriptional changes in oligodendrocytes (Fig. 7B). Notably, a separate single-nucleus transcriptomic study on chronic methamphetamine treatment in the mouse prefrontal cortex similarly reported that oligodendrocytes showed the most DEGs and observed evidence of myelin damage (Zeng, Yu et al. 2023).

Surprisingly, dopaminergic neurons displayed fewer DEGs in the sub-chronic treatment condition, both in comparison to the acute treatment group and relative to oligodendrocytes, astrocytes, and glutamatergic neurons in the sub-chronic group. We specifically examined the expression of genes involved in cholesterol metabolism within dopaminergic neurons. Strikingly, all examined genes showed a trend toward down-regulation, though not all changes reached statistical significance (Fig. 7C). The most significantly downregulated genes were *Ldlr* and *Msmo1* (Fig. 7D). This down-regulation of cholesterol metabolism genes may represent an adaptive response following the up-regulation observed in the acute treatment condition.

## Methods

### Mice

All animal procedures were conducted in accordance with institutional and national ethical guidelines and were approved by the Institutional Animal Care and Use Committee (IACUC) at National Institutes of Health. To facilitate accurate dissection of the VTA and SNc, *Slc6a3*-EGFP/*Rpl10a* (Dat bacTRAP) mice (Brichta, Shin et al. 2015) (Jackson Laboratory, Stock No. 012365) were used for identification of midbrain regions containing dopaminergic neurons. Dat bacTRAP mice exhibit EGFP fluorescence in dopaminergic neuronal cell bodies, enabling precise identification of the VTA and SNc under a fluorescence microscope.

RiboTag mice (Jackson Laboratory, Stock No. 029977)(Sanz, Yang et al. 2009) were crossed with Th-Cre mice (Jackson Laboratory, Stock No. 008601)(Savitt, Jang et al. 2005) to generate experimental animals. Mice heterozygous for both the RiboTag and Th-Cre alleles were used for all experiments. All animals were housed in a temperature- and humidity-controlled environment on a 12-hour light/dark cycle with ad libitum access to food and water. All efforts were made to minimize animal suffering and to reduce the number of animals used.

### Nuclear Isolation

Dat bacTRAP mice (10–16 weeks old) were used for all experiments. Mice received intraperitoneal injections of either saline or methamphetamine (3 mg/kg). For acute treatment, brain tissue was collected 3 hours post-injection. For repeated treatment, mice received daily injections for 7 consecutive days, and brains were harvested 24 hours after the final injection. Mice were anesthetized with isoflurane, and brains were quickly harvested in ice-cold artificial cerebrospinal fluid (ACSF; Ecocyte Bioscience, product code LRE-S-LSG-1000-1) bubbled with carbogen (95% O₂, 5% CO₂). Brains were sectioned at 1 mm thickness using a Zivic brain slicer matrix. Midbrain-containing slices were dissected under a fluorescent dissecting microscope, and only regions showing fluorescence—corresponding to the VTA and SNc—were collected. For each condition (3-hour and 7-day injections), 16 mice were used, with 8 animals per treatment group (4 males and 4 females for both saline and methamphetamine groups). Totally 32 animals were used.

The nuclei isolation protocol was performed as previously described(Lee, Rhodes et al. 2022, Lee, Zhang et al. 2023). Tissue from each mouse was homogenized in a 2ml Dounce Homogenizer containing 1 ml freshly prepared lysis buffer, applying 10 strokes with the A pestle followed by 10 strokes with the B pestle. The homogenate was filtered through a 40 µm cell strainer, transferred to a DNA low bind 2 mL microfuge tube and centrifuged at 300 × g for 5 min at 4 °C. The supernatant was removed, the pellet was gently resuspended in low sucrose buffer and centrifuged for another 5 min. The nuclei were resuspended in 500 µl resuspension buffer and loaded on top of 900 µl 1.8 M Sucrose Cushion Solution (Sigma, NUC-201). The sucrose gradient was centrifuged at 13,000 × g for 45 min at 4 °C. The supernatant was discarded; the nuclei of each sample were resuspended in 250 µl resuspension buffer. At this stage, samples from 4 mice (2 males and 2 females) within the same treatment group were pooled, resulting in a total volume of 1 ml per pooled sample. 5 ul of 5 mM DRAQ5 was added to each sample and filtered through a 35 µm cell strainer. Samples were processed on a Sony SH800 Cell Sorter with a 100 µm sorting chip. GFP^+^/DRAQ5^+^ nuclei were first collected to enrich dopaminergic neurons, then DRAQ5+ nuclei containing all cell types were collected and mixed into 1.5 ml centrifuge tubes containing 10 µl of the resuspend buffer. We collected about 15,000 nuclei.

### Single Nucleus Sequencing and Data Analysis

Using a Chromium Single Cell 3′ Library and Gel Bead Kit v3 (10X Genomics), nuclei were immediately loaded onto a Chromium Single Cell Processor (10X Genomics) for barcoding of RNA from single nuclei, and single cell libraries generated according to the manufacturer’s instructions. Resulting cDNA samples were run on an Agilent Bioanalyzer using the High Sensitivity DNA Chip as quality control to determine cDNA concentrations. The samples were combined and run on an Illumina HiSeq2500 with Read1 = 28 bp, Read2 = 91 bp, and indexRead = 8. Reads were aligned and assigned to Mouse Reference Genome Refdata Gex Mm10 2020 A using the CellRanger v7.1.0 pipeline (10X Genomics).

Sequencing data were analyzed using the R package Seurat version 5.1.0 (Stuart, Butler et al. 2019) following standard procedures(Butler, Hoffman et al. 2018). For each sample, Seurat objects were created from count matrices, retaining genes detected in at least 3 nuclei and nuclei with at least 200 detected genes. Additional filtering was performed to retain nuclei with more than 400 and fewer than 7,500 detected genes and fewer than 40,000 total UMI counts. This step excluded low-quality nuclei and potential doublets.

Data were log-normalized using the NormalizeData function in Seurat to correct for differences in sequencing depth and technical variation. 2 sets of data from saline- and methamphetamine-treated animals were integrated using Seurat’s Canonical Correlation Analysis (CCA)-based integration workflow (IntegrateLayers function, method = “CCAIntegration”) to minimize batch effects.

Clustering was performed using FindClusters, and clusters were visualized in two dimensions using UMAP. Clusters present only in a single sample were excluded. Cluster annotation was performed by examining expression of canonical marker genes and by comparison to published single-cell transcriptomic datasets.

For each major cell type, nuclei were grouped by treatment (saline or methamphetamine), and differential expression was performed using the FindMarkers function (Wilcoxon rank-sum test). Results were further filtered to include only genes expressed in more than 10% of nuclei in at least one group. All analyses were performed in Seurat v5.1.0, and visualization was performed using Seurat, ggplot2, and RColorBrewer.

### RNAscope, Imaging, and Data Analysis

RNAscope HiPlex *in situ* hybridization was performed using a two-round staining protocol to assess the expression of cholesterol metabolism–related genes in mouse brain sections, according to the manufacturer’s instructions (Advanced Cell Diagnostics, ACD). In the first round, probes for *Slc6a3, Ldlr*, and *Hmgcr* were hybridized and visualized, followed by fluorophore stripping and a second round with probes for *Th, Insig1*, and *Msmo1*.

All tissue sections were imaged using a Nikon Biopipeline widefield slide scanner with a Nikon plan-apochromat 40X/0.95 NA objective and Fusion sCMOS camera (2304×2304 pixels; 0.16 μm/pixel; 16-bit images). Fluorescent channels were configured for DAPI, Alexa 488, Alexa 555, and Alexa 647 with appropriate excitation and emission filters. Images were acquired as tiled regions with wavelength-specific shading correction, autofocusing, and z-stacks (20 μm range at 1 μm steps) at each ROI. Exposure times were set automatically to minimize saturation.

Image processing included stitching and flattening using an extended depth of focus algorithm, followed by deconvolution via the Richardson–Lucy algorithm (20 iterations, theoretical PSF). Image sets from subsequent staining rounds were registered to the initial round. Custom image processing scripts were implemented in MATLAB (R2019b) and ANTs (v2.2.0). Computational resources were provided by the NIH HPC Biowulf cluster (http://hpc.nih.gov).

RNA expression was quantified using Arivis Vision4D Pro software (v4.1.0, Build 16702). Regions of interest (ROIs) were defined by Slc6a3- or Th-positive cells. RNAscope puncta for each cholesterol-related gene were automatically counted within ROIs for each brain. Statistical comparisons between methamphetamine- and saline-treated groups were performed using GraphPad Prism.

### RNA Isolation from Dopaminergic Neurons Using *Th-cre:RiboTag* Mice

Mice were administered intraperitoneally (i.p.) with 3 mg/kg methamphetamine, amphetamine, methylphenidate, or saline. Four biological replicates were used per treatment group. RNA was specifically isolated from dopaminergic neurons using *Th-cre:RiboTag* mice, which express an HA-tagged ribosomal protein (Rpl22) in tyrosine hydroxylase (Th)-expressing cells. Three hours following injection, mice were euthanized, and brains were rapidly removed and sectioned to dissect out the VTA and SNc. Dissected tissues were flash-frozen in liquid nitrogen and stored at –80°C until use. For RNA isolation, frozen tissue was homogenized in ice-cold homogenization buffer (50 mM Tris-HCl, 100 mM KCl, 12 mM MgCl₂, 1% NP-40, 1 mM DTT, 200 U/mL RNase inhibitor, and protease inhibitors). Homogenates were clarified by centrifugation at 10,000 × g for 10 minutes at 4°C, and the supernatant was incubated with anti-HA magnetic beads (Thermo Fisher) for 4 hours at 4 °C with gentle rotation to immunoprecipitate HA-tagged ribosomes. Beads were washed three times with high-salt buffer (50 mM Tris-HCl, 300 mM KCl, 12 mM MgCl₂, 1% NP-40, 1 mM DTT, RNase inhibitor, and protease inhibitors). Ribosome-associated RNA was extracted from the beads using the RNeasy Mini Kit (Qiagen), including on-column DNase digestion to remove genomic DNA, according to the manufacturer’s protocol. RNA concentration and integrity were assessed using an Agilent TapeStation, and only samples with an RNA integrity number (RIN) greater than 7 were used for downstream analyses.

### qPCR Analysis of Cholesterol Metabolism Genes in Response to Methamphetamine Administration

For quantitative PCR (qPCR) analysis, ribosome-associated RNA was first amplified using the SuperScript™ IV Single Cell/Low Input cDNA PreAmp Kit (Thermo Fisher Scientific) according to the manufacturer’s instructions. Amplified cDNA was then subjected to qPCR using TaqMan™ Gene Expression Assays (Thermo Fisher Scientific) specific for cholesterol metabolism-related genes. Reactions were prepared with TaqMan™ Fast Advanced Master Mix (Thermo Fisher Scientific) and run on a ViiA 7 Real-Time PCR System (Applied Biosystems). Each sample was analyzed in technical triplicate. Relative gene expression was calculated using the ΔCT (delta CT) method, with *Gapdh* serving as the endogenous control.

### Bulk RNA Sequencing and Data Analysis

Bulk RNA-seq libraries were prepared using the Clontech SMARTer Ultra Low Input RNA Kit v4 (Takara Bio) according to the manufacturer’s instructions. Sequencing libraries were generated with the Nextera™ DNA Flex Library Prep Kit (Illumina) and subjected to paired-end sequencing (2 × 150 bp) on an Illumina HiSeq 4000 platform. Raw sequencing reads were assessed for quality using FastQC, and adapters and low-quality bases were trimmed using Cutadapt. Trimmed reads were aligned to the mouse reference genome (mm10) and annotated transcripts using STAR. Gene expression quantification was performed for all samples using RSEM. Differential gene expression analysis was carried out using the DESeq2 package in R. Pathway and gene ontology analyses were performed using Metascape.

## Discussion

In this study, we performed snRNA-seq to demonstrate that acute methamphetamine exposure leads to robust and selective up-regulation of cholesterol metabolism genes in midbrain dopaminergic neurons. Our data show that this transcriptional response is broad, encompassing key enzymes, transcription factors, and regulatory genes involved in cholesterol synthesis, uptake, and homeostasis. While astrocytes also exhibit increased expression of some cholesterol metabolism genes, fewer DEGs in the pathway were detected when compared to dopaminergic neurons. Other major midbrain cell types, including oligodendrocytes, GABAergic, and glutamatergic neurons, display minimal changes in cholesterol-related gene expression following acute methamphetamine administration. Notably, we find that amphetamine induces a similar up-regulation of cholesterol metabolism genes in dopaminergic neurons, whereas methylphenidate does not, highlighting divergent effects among commonly used psychostimulants. The selective transcriptional activation of cholesterol metabolism genes in response to amphetamine-derivatives, but not methylphenidate, further underscores the specificity of this effect. Whereas both amphetamine and methamphetamine interact with DAT to increase extracellular dopamine, they also activate intracellular signaling pathways directly through their actions on TAAR1 (Underhill, Hullihen et al. 2021). In contrast, methylphenidate primarily blocks reuptake with less effect on dopamine release (Challman and Lipsky 2000, Faraone 2018). Together, these results suggest that while amphetamine and methamphetamine can affect dopaminergic signaling through their intracellular actions, methylphenidate, which blocks DAT but does not significantly enter the neurons, exerts its action through distinct mechanisms.

The mechanisms underlying the robust up-regulation of cholesterol metabolism genes are intriguing. Work from our lab and others has shown that amphetamines can activate the RhoA–ROCK signaling pathway in dopaminergic neurons (Narita, Takagi et al. 2003, Wheeler, Underhill et al. 2015). More recently it has been shown that amphetamine can enter cells and activate the intracellular trace amine-associated receptor 1 (TAAR1), which is coupled to the G protein G₁₃. TAAR1 signaling through G₁₃ activates RhoA at or near the endoplasmic reticulum (Underhill, Hullihen et al. 2021). RhoA activation subsequently triggers endocytosis of DAT and a neuronal glutamate transporter, excitatory amino acid transporter 3 (EAAT3) (Underhill, Wheeler et al. 2014, Wheeler, Underhill et al. 2015, Underhill, Colt and Amara 2020). Endocytosis from cholesterol-enriched lipid rafts can lead to redistribution of cholesterol from plasma membrane to endosomes (Yue and Xu 2015, Mu, Li et al. 2017, Ikonen and Zhou 2021), which may result in local depletion of cholesterol at the plasma membrane. Such depletion is a well-established trigger for activation of the SREBP pathway, which transcriptionally upregulates cholesterol biosynthesis genes. In addition, independent of its role in endocytosis, RhoA–ROCK signaling has been shown to directly promote SREBP processing and nuclear translocation(Lin, Zeng et al. 2003). Together, these findings suggest that amphetamine-induced RhoA activation may drive SREBP-mediated transcription of cholesterol biosynthesis genes through both membrane remodeling and direct intracellular signaling mechanisms.

The functional significance of cholesterol changes in dopaminergic neurons is especially noteworthy. Dopaminergic neurons are among the most metabolically active cells in the brain, and their function is tightly linked to the activity of DAT, which regulates extracellular dopamine levels by reuptake. Cholesterol is a critical component of neuronal membranes and directly modulates DAT conformation and activity. Previous work, including studies by our group and others, has shown that cholesterol maintains the outward-facing conformation of DAT (Hong and Amara 2010, Wang, Neel et al. 2023), and that reducing membrane cholesterol with statins significantly decreases dopamine uptake. Thus, the up-regulation of cholesterol metabolism genes observed after psychostimulant exposure may serve to support DAT function and synaptic homeostasis during periods of intense dopaminergic activity, potentially influencing both dopamine uptake, efflux and neuronal excitability in response to drugs like methamphetamine and amphetamine. Cholesterol is essential for synaptic vesicle formation, neurotransmitter release, and membrane repair—all processes that are acutely stressed during heightened dopamine turnover. By rapidly enhancing cholesterol synthesis, dopaminergic neurons may support increased synaptic activity and maintain membrane integrity in the face of pharmacological stress.

Importantly, our study showed that *Hmgcr*—gene encoding the rate-limiting enzyme in cholesterol synthesis and the target of statins—is highly enriched in dopaminergic neurons and is upregulated after methamphetamine exposure. This observation has significant implications, as statins are widely prescribed cholesterol-lowering drugs that cross the blood-brain barrier to varying degrees. Previous studies suggest that statins may exert neuroprotective effects on dopaminergic neurons, providing therapeutic benefits for Parkinson’s disease (Selley 2005, Ghosh, Roy et al. 2009, Al-Kuraishy, Fahad et al. 2024). Our findings suggest that statins might affect dopaminergic neuron function potentially also in the context of psychostimulant exposure.

Beyond neuroprotection, our dynamic gene expression data imply that statins could influence vulnerability to psychostimulant addiction and relapse. Supporting this idea, studies have shown that simvastatin can reverse cocaine-induced changes in lipid profiles in the nucleus acumens and block the reinstatement of cocaine-conditioned place preference (Xu, He et al. 2021). Additional work has demonstrated that statins reduce cocaine and nicotine addiction behavior in rats (Chauvet, Nicolas et al. 2016) and prevent morphine-induced tolerance and dependence in mice (Pajohanfar, Mohebbi et al. 2017). These findings, together with our results, suggest that cholesterol metabolism may modulate addictive behaviors and the brain’s response to psychostimulants. Interestingly, psychostimulants and statins are widely used in the population, and thus further exploration of the potential interaction of these medications are needed.

Peripheral cholesterol cannot readily cross the blood-brain barrier, and traditional models held that neurons depend on astrocyte-derived cholesterol supplied via ApoE-containing particles (Nieweg, Schaller and Pfrieger 2009, Li, Zhang and Liu 2022). However, our data show that dopaminergic neurons themselves are highly enriched in the expression of genes required for cholesterol biosynthesis and exhibit robust up-regulation following acute methamphetamine exposure. The up-regulation of both Bloch (Dhcr7) and Kandutsch-Russell (*Dhcr24*) pathway genes suggests that dopaminergic neurons possess strong intrinsic cholesterol biosynthetic capacity. We also observed an increased *Apoe* expression in astrocytes, which suggests the possibility of crosstalk and coordinated lipid homeostasis. Whether the response is primarily cell-autonomous or a coordinated effort between neurons and astrocytes remains to be fully elucidated. Nonetheless, our findings support the conclusion that dopaminergic neurons are likely more self-sufficient in cholesterol synthesis than previously appreciated.

## Conclusions

In summary, our findings reveal that acute exposure to methamphetamine and amphetamine induces a rapid and robust up-regulation of cholesterol metabolism genes in midbrain dopaminergic neurons. This adaptive metabolic response is highly cell-type– specific, drug-specific, and dynamic over time, offering new insights into the role of cholesterol in dopaminergic neuron function and addiction. Furthermore, our study demonstrates that within the midbrain, dopaminergic neurons are the most enriched for genes associated with cholesterol metabolism, challenging the prevailing view that neurons are merely passive recipients of astrocyte-derived cholesterol.

---

**Supplementary Figure 1.**
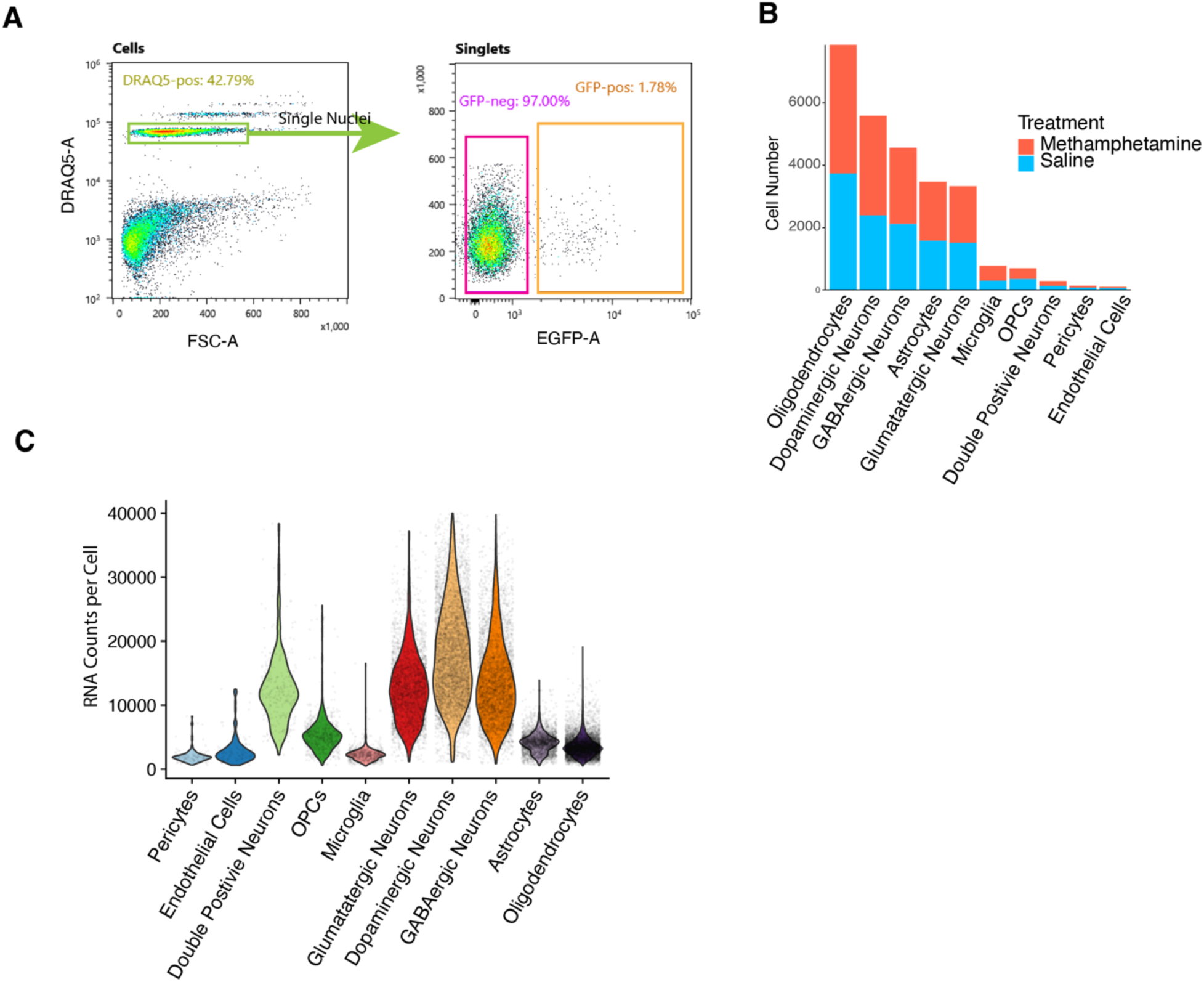
Overview of single-nucleus RNA-seq data. (A) FACS gating strategy for nuclei collection. DRAQ5⁺ single nuclei were identified and subdivided into GFP⁺ nuclei (from dopaminergic neurons) and GFP⁻ nuclei. GFP⁺ nuclei were collected first, followed by GFP⁻ nuclei, resulting in a mixed nuclei population enriched for dopaminergic neurons. (B) Numbers of nuclei from each cell type passing quality control in the RNA-seq dataset. OPCs, oligodendrocyte precursor cells. (C) Violin plot showing RNA counts per cell for each cell type.

**Supplementary Figure 2.**
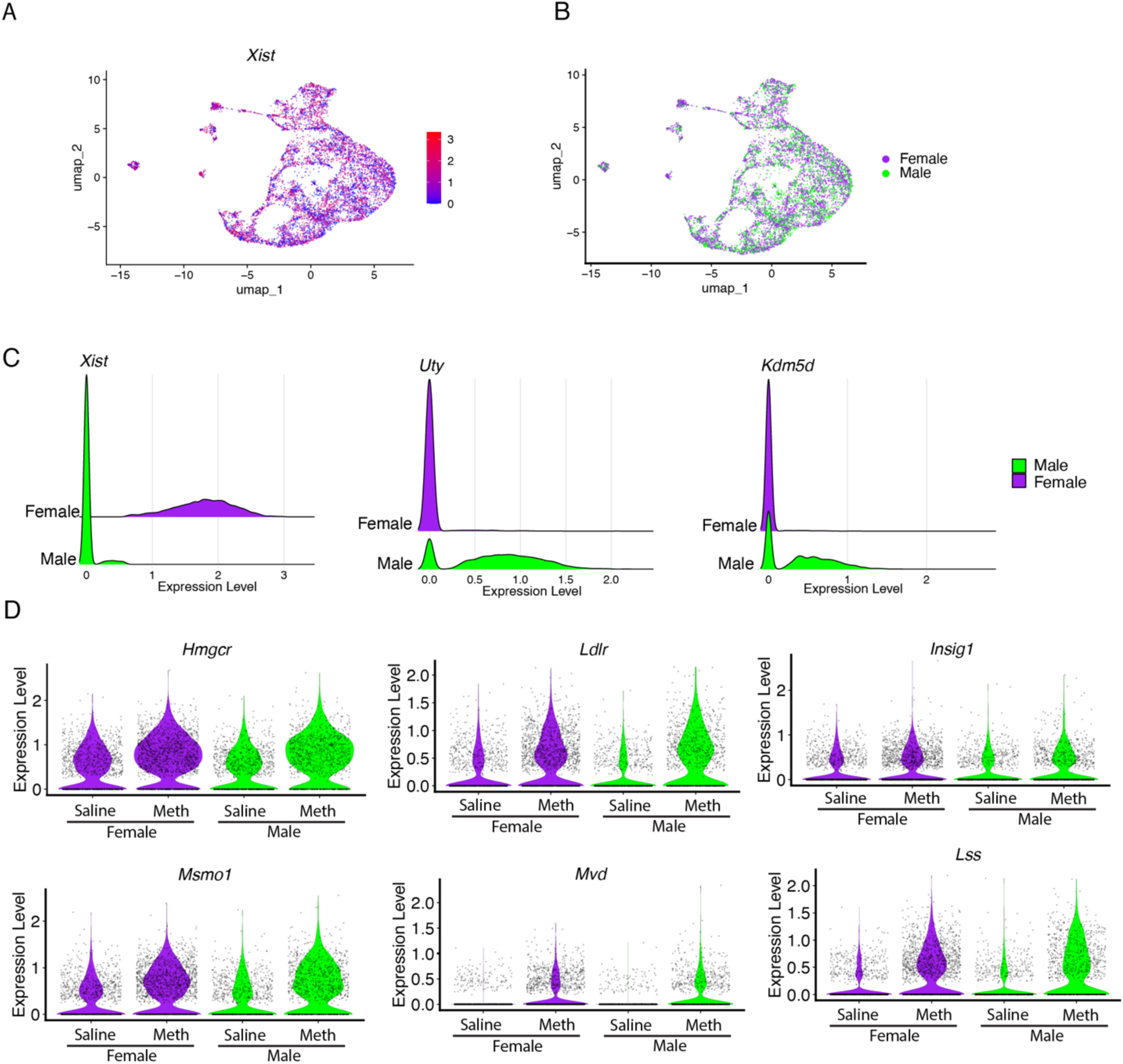
Expression of cholesterol metabolism genes in male and female dopaminergic neurons. (A) UMAP plot of dopaminergic neurons showing the distribution of *Xist* expression (female-specific marker). (B) Separation of male and female dopaminergic neurons based on *Xist* expression. (C) Ridge plots displaying expression of the female-specific gene *Xist* and male-specific genes *Uty* and *Kdm5d* in male and female cells. (D) Violin plots showing expression levels of selected differentially expressed genes (DEGs) associated with cholesterol metabolism in male and female dopaminergic neurons following acute methamphetamine treatment.

**Supplementary Figure 3.**
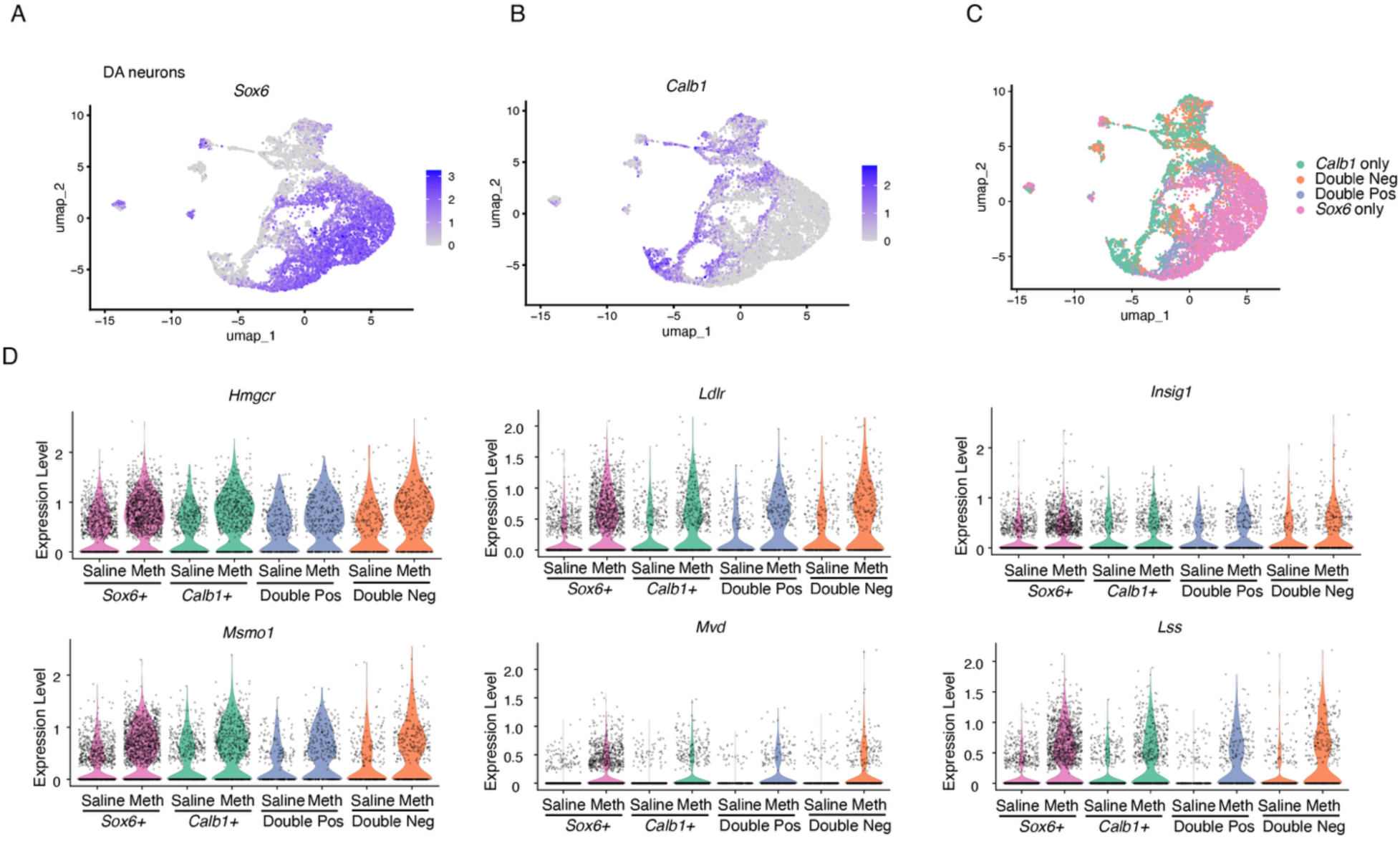
Expression of cholesterol metabolism–related genes in VTA and SNc dopaminergic neuron subpopulations. (A) UMAP plot showing expression of *Sox6*, a marker enriched in SNc dopaminergic neurons. (B) UMAP plot showing expression of *Calb1*, a marker enriched in VTA dopaminergic neurons. (C) UMAP plot illustrating dopaminergic neuron subpopulations divided into *Calb1*-only, *Sox6*-only, double-negative, and double-positive groups based on marker expression. (D) Violin plots displaying expression levels of selected differentially expressed genes (DEGs) related to cholesterol metabolism in the four location-associated dopaminergic neuron subpopulations following acute methamphetamine treatment.

**Supplementary Figure 4.**
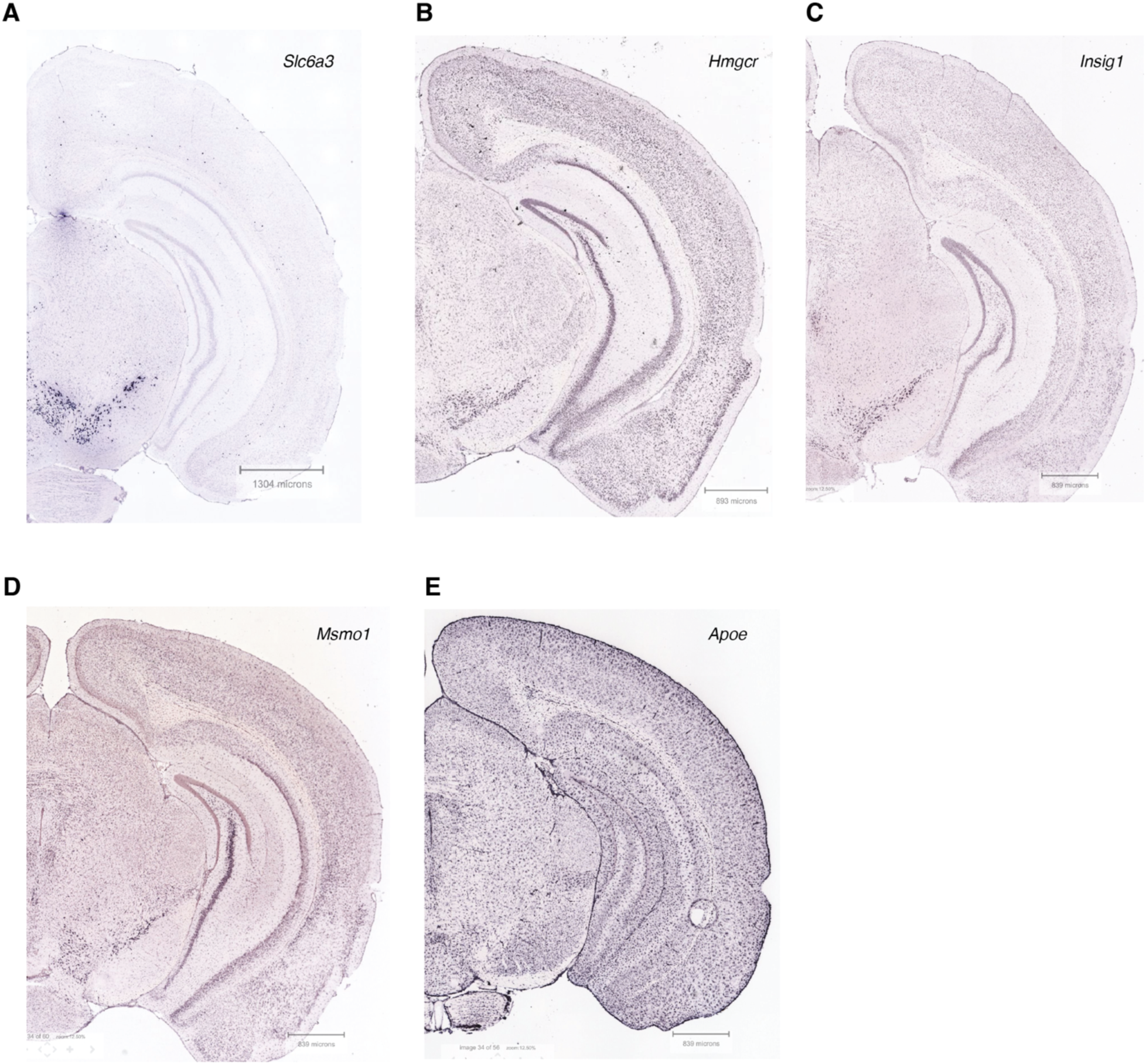
*In situ* hybridization images from the Allen Brain Atlas showing distribution of genes involved in cholesterol metabolism. (A)The expression pattern of dopaminergic marker gene *Slc6a3*. (B-E) The expression patterns of genes involved in cholesterol metabolism.

**Supplementary Figure 5.**
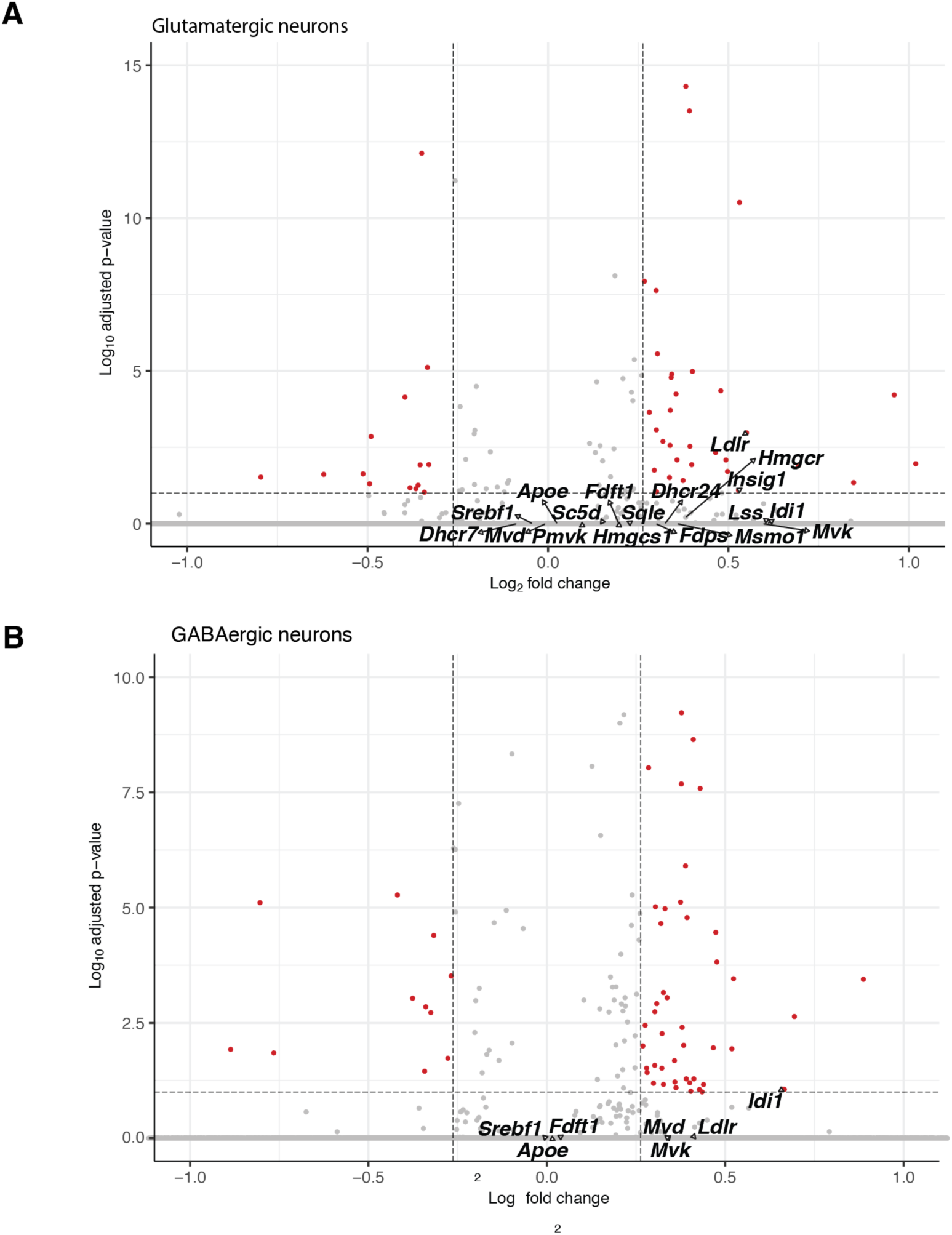
Expression of cholesterol metabolism–associated genes in non-dopaminergic neurons. (A, B) Volcano plots displaying adjusted p-values and log₂ fold changes for cholesterol metabolism–related genes in glutamatergic neurons (A) and GABAergic neurons (B) following acute saline or methamphetamine treatment. Differentially expressed genes were identified using a cutoff of FDR < 0.1 and absolute fold change > 1.2.

